# Tensor Factorization-based Prediction with an Application to Estimating the Risk of Chronic Diseases

**DOI:** 10.1101/810556

**Authors:** Haolin Wang, Qingpeng Zhang, Frank Youhua Chen, Eman Yee Man Leung, Eliza Lai Yi Wong, Eng-Kiong Yeoh

## Abstract

Tensor factorization has emerged as a powerful method to address the challenges of high dimensionality regarding disease development and comorbidity. Chronic diseases have a high likelihood to co-occur, making patients suffering from one chronic disease to have an elevated risk for the other diseases in the course of aging. Individualized prediction of chronic diseases can help patients prevent new diseases and reduce the healthcare costs. Despite rich results of risk assessment models for chronic diseases, individualized risk prediction considering the complex mechanisms of disease development and comorbidity remains to be under-researched. This research aims to develop tensor factorization-based machine learning models to predict the onset of new chronic diseases for individual patients through incorporating the comorbidity patterns with the clinical and sequential factors revealed in the electronic health records (EHR) data. We propose two tensor factorization-based methods to incorporate the clinical and sequential factors to reveal the latent patterns of co-occurring chronic diseases. The efficacy of the proposed methods was validated through predicting the onset of new chronic diseases for individual patients using the EHR data for 23 years from a major hospital in Hong Kong. The proposed methods consistently outperform benchmark predictive models. The top 10 predictions of new chronic diseases have approximately 60% recall. Tensor factorization is an appropriate method for predicting the onset of chronic diseases at the individual level. The proposed predictive models could inform proactive health management programs for at-risk patients with different chronic conditions at discharge.

**Author summary:** The existing risk assessment models mainly focused on the prediction of single diseases in the population base. Chronic disease risk prediction considering the complex mechanisms of disease development and comorbidity is under-researched. To support and inform clinical decision making for healthcare professionals in the aging society, this study provides an innovative approach to mapping an interconnected web of chronic illnesses and investigated the performance of chronic disease prediction using 2 years’ worth of patient assessment records and 23 years’ admission history data from a major hospital in Hong Kong. We proposed matrix and tensor-based methods to represent the high-order interrelations of patients, chronic diseases and additional features, which can reveal the latent patterns of co-occurring chronic diseases to enable more effective prediction. The proposed methods exhibit state-of-the-art performance in predicting the onset of new chronic diseases for individual patients.

## Introduction

Tensor factorization, the high order extension of the two-dimensional matrix factorization, has emerged as a promising method to address the challenges regarding the high dimensionality of the EHR data with good interpretability and scalability [1, 2]. Tensor factorization has been widely used in recommender systems, social network analysis, process monitoring etc. [3–5]. Recently, tensor-based models have also been applied to healthcare problems, including phenotype generation, medical information retrieval, image-based diagnosis, and precision medicine [6–11].

Compared with traditional machine learning methods, tensor-based models have the unique advantages of having: (a) the capability to utilize multi-aspect features in multiple dimensions; (b) the versatility in incorporating domain knowledge from physicians or knowledge bases in medicine; and (c) the capability to solve the sparsity problem, a major challenge for many data mining tasks, particularly for an EHR dataset. These advantages make tensor factorizations a promising modeling approach to disease prediction using the EHR data, which usually have high dimensionality and sparsity, and needs domain expertise to ensure model validity [1, 12, 13].

Chronic diseases are a major cause of morbidity and mortality worldwide [14]. As of 2012, approximately half of all adults in the US had one or more chronic health conditions, and one in four adults had at least two chronic conditions [15]. The reasons for the rapid rise in chronic illness include the population aging, longer life expectancies due to improvements in medical care, and advances in diagnostic technology and treatment options for many chronic diseases. Among older adults in the US, 77% of them have at least two chronic illnesses [16] and 43% of Medicare beneficiaries have three or more [17]. For instance, chronic conditions such as hypertension, heart diseases, diabetes and chronic obstructive pulmonary disease (COPD) have a high likelihood to co-occur, making patient suffering from any one of these five chronic conditions to have an elevated risk for the other four conditions in the course of aging [18–20]. In addition, even acute conditions that are frequently considered as causes for hospitalization among the elderly, such as sepsis, peritonitis or fall, they could, nonetheless, be a manifestation of the underlying chronic conditions. It has also been recognized that early risk identification can facilitate early prevention/disease management in the community, thereby reducing the number of people suffering from chronic diseases and their acute presentations [21]. However, it is not until the widespread adoption of Electronic Health Records (EHRs) could predictive analytics be applied to shed light on the evolution and comorbidity of chronic diseases [22–24].

Harnessing the EHRs data to predict diseases has emerged as an important topic for medical informatics and precision medicine [25–27]. Predictive models based on EHR data can help physicians assess the future risk of an individual for a certain chronic condition or multiple conditions and identify the patients who are likely to acquire new diseases. Moreover, predictive models enable the comparison of benefits and the costs/risks of alternative treatments and prevention strategies, and allows for personalized disease management for individuals [28, 29].

Existing research on chronic diseases prediction, and subsequent prevention and management, has mainly focused on developing regression models to estimate the clinical risk factors and genomic variables, such as biomarkers, medical history, family history and genealogy records, and demographics [28, 30]. For instance, QDScore was proposed as the first diabetes prediction (regression-based) algorithm for type-2 diabetes [31]. In another study, logistic regression-based DiaRem score was proposed to predict the readmission of type-2 diabetes after Roux-en-Y gastric bypass surgery [32]. Similar research on developing regression models for diabetes prediction is rich and has been validated with various datasets [33–35]. In addition to diabetes, successful regression-based risk assessment models exist for other chronic diseases, like cardiovascular and chronic kidney disease [36–38]. Refer to a recent review for details [12].

During the past decade, machine learning models have been recognized as an effective method for chronic disease predictions. For instance, Himes et al. developed a Bayesian network model to predict COPD in asthma patients, and demonstrated the good accuracy of the model using 15 years of the EHR data [39]. Kurosaki et al. constructed decision tree models that can identify patients at a high risk of hepatocellular carcinoma development among different datasets [40, 41]. In another study, artificial neural networks and C5.0 classifiers were merged with decision trees to form a hybrid model to predict type-1 diabetes mellitus [42]. A recent study evaluated the performance of three classic models (naïve Bayesian classifier, Bayesian network, and support vector machines) in leveraging daily self-monitoring reports to predict asthma exacerbation and demonstrated the potential of machine learning models in providing personalized monitoring decision support [28].

Despite the rich results of risk assessment models for chronic diseases, the existing literature mainly focused on the prediction of single diseases in the population base, but not the individualized risk prediction considering the complex mechanisms of disease development and comorbidity, which usually have a high dimensionality [12, 43].

This research proposes two third-order tensor factorization-based models to predict the onset of new chronic diseases for individual patients by uncovering the latent comorbidity patterns of chronic diseases. Both models are extended from a second-order patient-disease matrix to a third-order tensor. One model incorporates the clinical assessment factors as the third dimension, whereas the other model incorporates the sequential factors as the third dimension. In developing new models for risk analytics related to chronic diseases, the current research aims to develop predictive and prescriptive models to improve the accuracy of risk assessment of underlying and co-occurring chronic conditions and has the potential to facilitate better post-discharge individualized care planning and patient education concerning one’s future risk of chronic and acute conditions given the patients’ chronic conditions.

This study provides an innovative approach to mapping an interconnected web of chronic illnesses and therefore contributes to the research in using advanced machine learning techniques to support and inform clinical decision making for healthcare professionals in our aging society. By identifying the unique health trajectories of patients, the proposed tensor factorization-based predictive models can predict the onset of new chronic diseases at the individual patient level. Compared with traditional regression and machine learning models, the proposed tensor factorization-based model is not only more accurate but is also more sensible and actionable for healthcare professionals to develop personalized prevention programs to reduce the chance of acquiring new chronic diseases and getting hospitalized.

## Methods

### Data

Two datasets were obtained from the internal medicine department of a major hospital in Hong Kong. The first dataset, admission history dataset, comprises 23 years (1993 to 2015) of diagnoses records from 20,070 patients. Another dataset for 2014 and 2015 (patient assessment dataset) contains the patient assessment information (e.g. heart rate, blood pressure, smoking). The two datasets contain 39 prevalent chronic diseases in Hong Kong. The diagnoses were encoded with the International Classification of Diseases (ICD-10) standard. To classify diseases, the commonly used Clinical Classifications Software and Chronic Condition Indicator [44] were adapted. Noting that acute myocardial infarction (most of which caused by coronary artery disease) caused by chronic conditions is included. The data were manually screened to ensure the encoding was consistent. Ten most common chronic diseases with occurrence and ICD-10 codes are shown in Table 1.

**Table 1.** Chronic disease classification.

| Disease Category | ICD-10 Code | # of patients |
| --- | --- | --- |
| Essential hypertension | I10 | 3417 |
| Chronic kidney disease | N18.5, N18.9, Z94.0, Z99.2 | 3238 |
| Diabetes mellitus without complication | E10.9, E11.9 | 1998 |
| Chronic obstructive pulmonary disease and bronchiectasis | J43.9, J44.9 | 1944 |
| Coronary atherosclerosis and other heart disease | I20.0, I20.9, I24.8, I25.2, I25.5, I25.9 | 1908 |
| Acute myocardial infarction | I21.4 | 1283 |
| Disorders of lipid metabolism | E78.0, E78.1, E78.2, E78.4 | 931 |
| Secondary malignancies | C77.0, C77.1, C77.2, C77.8, C77.9, C78.2, C78.6, C78.7, C79.9 | 632 |
| Other endocrine disorders | E16.0, E16.1, E16.2, E20.9, E21.0, E21.3, E22.0, E22.2, E23.0, E23.2, E23.6, E23.7, E24.2, E24.9, E27.1, E27.2, E27.8 | 511 |
| Deficiency and other anemia | D50.0, D56.0, D56.1, D56.3, D56.9, D58.0, D58.9, D59.1, D59.4, D59.5, D59.6, D59.9, D60.9, D61.3, D61.9 | 456 |

### Preliminaries on Matrix and Tensor Factorizations

Matrix factorization decomposes a matrix into the product of matrices [45]. Tensor factorization is the high order extension of matrix factorization and enables the modeling of heterogeneous and multidimensional data. Matrix and tensor factorizations can extract the latent components to enhance data mining tasks [1]. The most widely used tensor factorization methods are CANDECOMP/PARAFAC (CP) factorization and Tucker factorization [5]. The Tucker factorization decomposes a tensor into a core tensor multiplied by a matrix along each mode. Meanwhile, CP factorization decomposes the tensor into the sum of rank one tensors. CP factorization is also a special case of Tucker factorization, in which the core is superdiagonal. CP factorization is unique under mild assumptions, making it suitable to uncover and interpret the actual latent factors because no equivalent rotated factorization yields the same fit [1]. The rest of this paper adopts CP factorization to perform prediction.

To illustrate matrix and tensor factorizations, we present the factorizations of a second-order matrix ***χ*** ∈ ***R***^*I*×*J*^ and a third-order tensor ***χ*** ∈ ***R***^*I*×*J*×*K*^ in Fig 1. The matrix and the tensor can be expressed as the following Eq (1) and Eq (2), respectively:

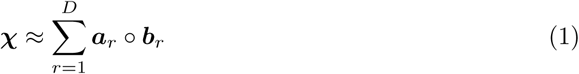

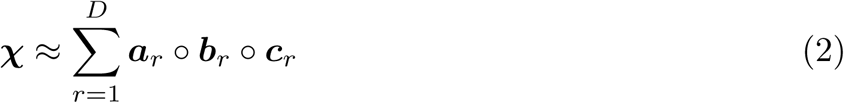

where ∘ denotes the outer product.

**Fig 1.**
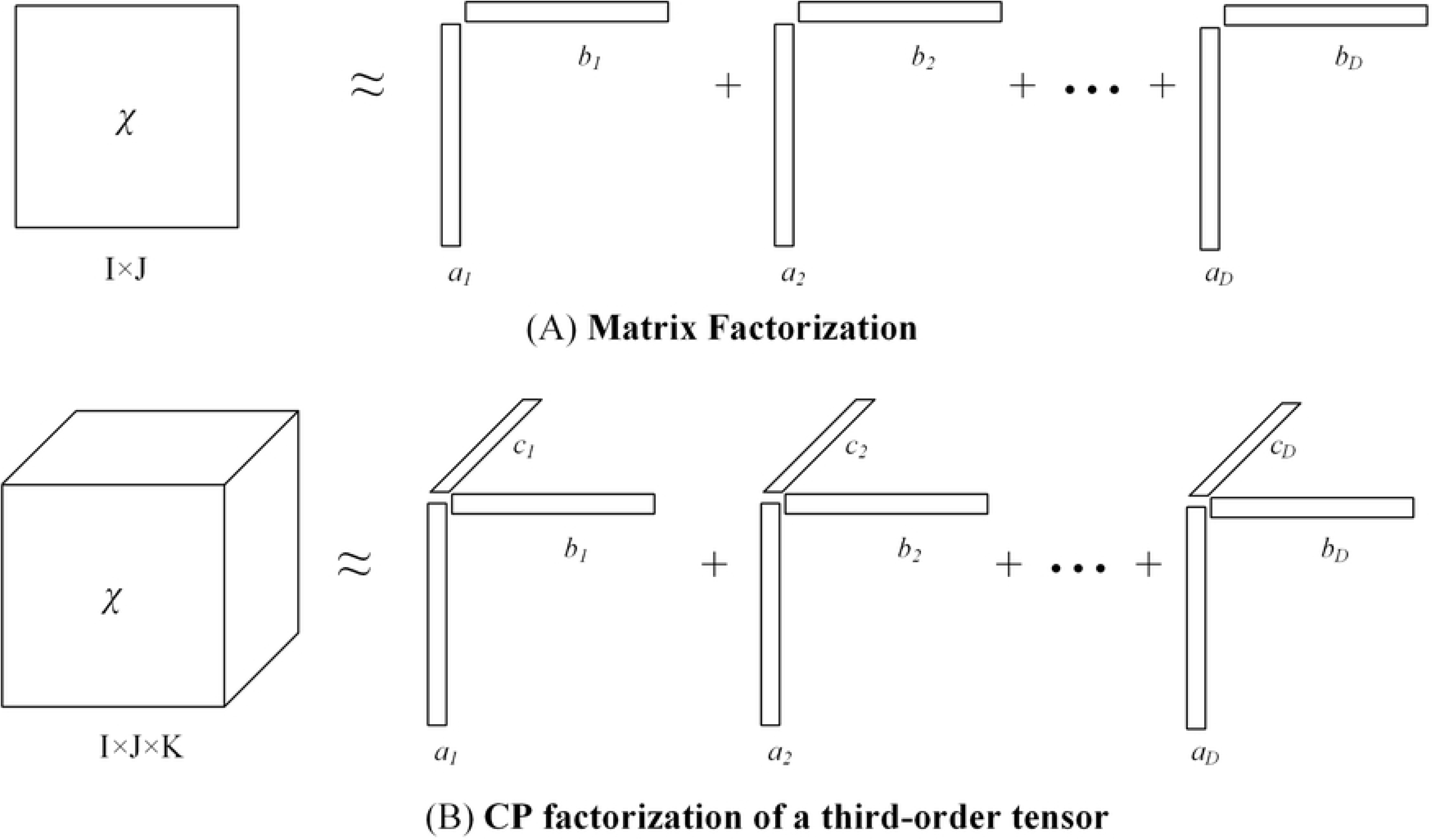
Illustration of matrix and third-order tensor factorizations.

Each data entry in the second-order matrix (*x*_*ij*_) could be interpreted as the inner product of two latent feature vectors, as shown by Eq (3). Similarly, each data entry in the third-order tensor (*x*_*ijk*_) could be interpreted as the inner product of three latent feature vectors, as shown by Eq (4). Such factorization is highly interpretable for multidimensional data mining applications, as we can interpret the decomposed components as high-order grouping patterns [1].

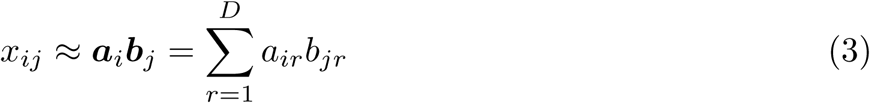

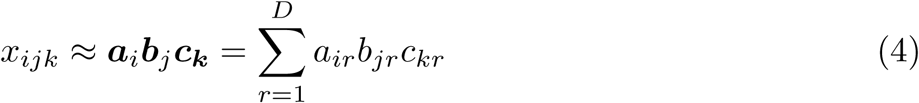

Many matrix and tensor factorization methods have different assumptions regarding factors and the underlying structures. Particularly, nonnegative matrix and tensor factorizations, both of which incorporate nonnegative constraints, have proven to be successful in many applications [46]. Such nonnegative constraints are suitable for this study because the values of EHR data entries are mostly nonnegative. The most popular cost functions are (a) the least squares error that corresponds to an assumption of normal independently and identically distributed noise, and (b) the Kullback-Leibler (KL) divergence that corresponds to maximum likelihood estimation under an independent Poisson assumption [3, 47]. As the count of admissions is used to construct tensors in the predictive models, we adopt the tensor factorization methods using generalized KL divergence and the multiplicative update rules [48, 49]. Refer to a recent review for the details of tensor factorizations [1].

### Matrix and Tensor Factorizations for Chronic Disease Prediction

In this study, we use matrices to represent the two-dimensional relationship between patients and diseases. Third-order tensors are constructed to represent the high-dimensional interrelations among patients, chronic diseases, and additional features. The matrices and tensors constructed by the EHR data are usually sparse with numerous “Nil” values. For example, there are many possible chronic diseases, and a patient usually has a few of them upon discharge. There is no value for the rest of the diseases, meaning that the patient has not acquired these diseases yet. Estimating which disease will be likely acquired by this patient is difficult. Matrix and tensor factorizations have been demonstrated to be effective estimating the values of such “Nil” data through exploring the latent grouping patterns in the observed tensors [50]. In the context of disease prediction, we can use matrix/tensor factorization to extract the latent grouping patterns of patients and diseases and reconstruct the observed matrix/tensor with factorized ones. Then, the updated values (in the reconstructed matrix/tensor) for those “Nil” entries can be used to estimate the risks of acquiring different diseases.

### Nonnegative Matrix Factorization (NMF)

Initially, as a baseline method to illustrate the factorization approach for prediction, the NMF methods are adopted to characterize the patients and chronic diseases by the vectors of factors inferred from the ⟨*patients, diseases*⟩ matrix in Fig 2(a). The data entries in the matrix are binary values to indicate whether the patient has the corresponding chronic disease.

**Fig 2.**
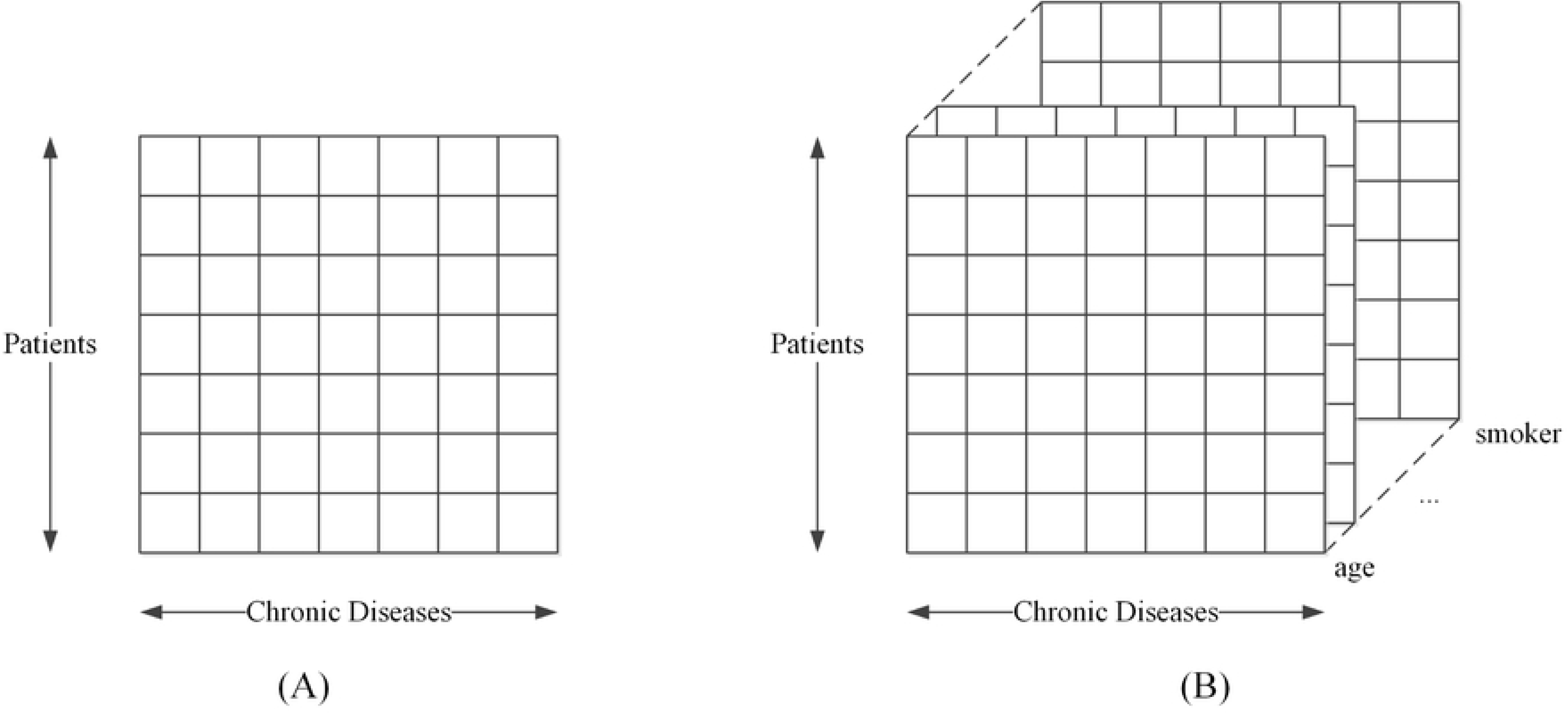
Illustration of (a) the matrix representation of the relations of patients and diseases and (b) the tensor representation of the ternary interrelations of patients, diseases and clinical attributes.

We decompose the matrix as shown in Fig 1(a) and obtain the feature factors of patients (*p*_*i*_) and chronic diseases (*d*_*j*_). The predicted risk score for patient *i* to acquire chronic disease *j* is the inner product of extracted latent feature vectors as

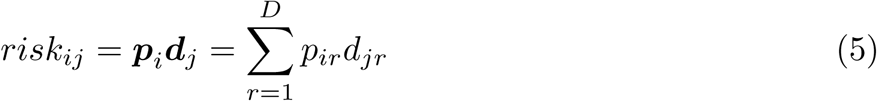

### Nonnegative Tensor Factorization with Clinical information (NTF-C)

Matrix-based data mining methods lack the capability to capture the characteristics and patterns in multi-aspect data. Medical research has recognized that clinical attributes could indicate the patients’ various health trajectories, which could lead to acquiring different diseases in the future [51]. For instance, hypertension could lead to several chronic diseases for the patient, given the specific clinical attributes (e.g. certain symptoms) he/she has at present. Patients who smoke are more likely to acquire respiratory disease like COPD, whereas patients with irregular heartbeat could be on the path to cardiovascular disease. Tensor factorizations provide a powerful framework to model such multi-aspect data by explicitly exploiting the multi-aspect structure to identify the latent clusters of data [1]. We extend the second-order NMF model into a third-order nonnegative tensor factorization (NTF) method to capture the clinical attributes.

To model the multi-aspect interrelations among patients, chronic diseases, and clinical attributes, we construct a third-order observed tensor ***χ*** (as shown in Fig 2(b)), in which *x*_*ijk*_ represents the frequency of admissions that comprise the corresponding ternary interrelations of ⟨*patient i, chronic disease j, clinical attribute k*⟩. Our model, named Nonnegative Tensor Factorization with Clinical information (NTF-C), decomposes observed tensor ***χ*** to the sum of rank-one tensors as illustrated in Fig 1(b). *x*_*ijk*_ can then be estimated through the inner product of three latent feature vectors.

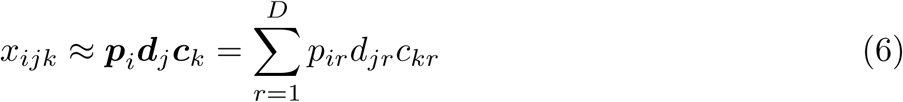

where *p*_*i*_, *d*_*j*_, and *c*_*k*_ are the feature vectors of patients, chronic diseases, and clinical attributes, respectively.

Then, ***R*** is defined as the reconstructed tensor. Specifically, ***R*** is obtained by taking the outer product of the factorized components. The values of data entries *r*_*ijk*_ in the reconstructed tensor are the estimated values for *x*_*ijk*_ obtained by Eq (6). This reconstructed tensor ***R*** is not equal to the observed tensor ***χ***; instead, ***R*** is a low-rank approximation of ***χ***. This reconstruction process can capture the latent high-dimensional grouping patterns of ***χ***. Thus, ***R*** provides evidence for predicting the values of the entries that are “Nil” in the observed tensor ***χ***. The risk factors are determined by the values of entries in ***R***. If the entry is “Nil” in the observed tensor ***χ***, but positive in the reconstructed tensor ***R***, it indicates a risk of acquiring this disease for the corresponding patient. In the tensor, a tube represents risk factors embedded in different clinical attributes for patient *i* and disease *j*. This set of risk factors is integrated across clinical attributes to generate a risk score for patient *i* in acquiring disease *j* as follows:

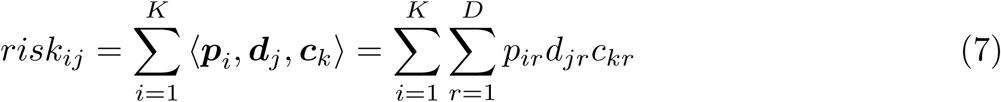

### Nonnegative Tensor Factorization with Sequential Information (NTF-S)

The chronic diseases of patients evolve over time with a complicated comorbidity relationship among different diseases [20]. Acquiring one chronic disease may lead to the risk of subsequently acquiring another chronic disease. Another third-order tensor-based model is proposed to model the sequence of chronic diseases. As shown in Fig 3, the sequence of diagnoses is incorporated into the model with a two-slice approach. Each admission of patient *i* due to an existing chronic disease *j* is recorded in slice *T* by incrementing 1 to ***χ***_*ijT*_. If another chronic disease *m* is also present, it is also recorded in slice *T* by incrementing 1 to ***χ***_*imT*_. The occurrence of a new chronic disease *n* will be recorded in slice *T* + 1 by incrementing 1 to ***χ***_*in*(*T* +1)_. By default, the diseases diagnosed in the previous visit of a patient are considered as existing diseases (recorded in slice *T*). In this way, the comorbidity information and the sequential pattern of these diseases are both represented.

**Fig 3.**
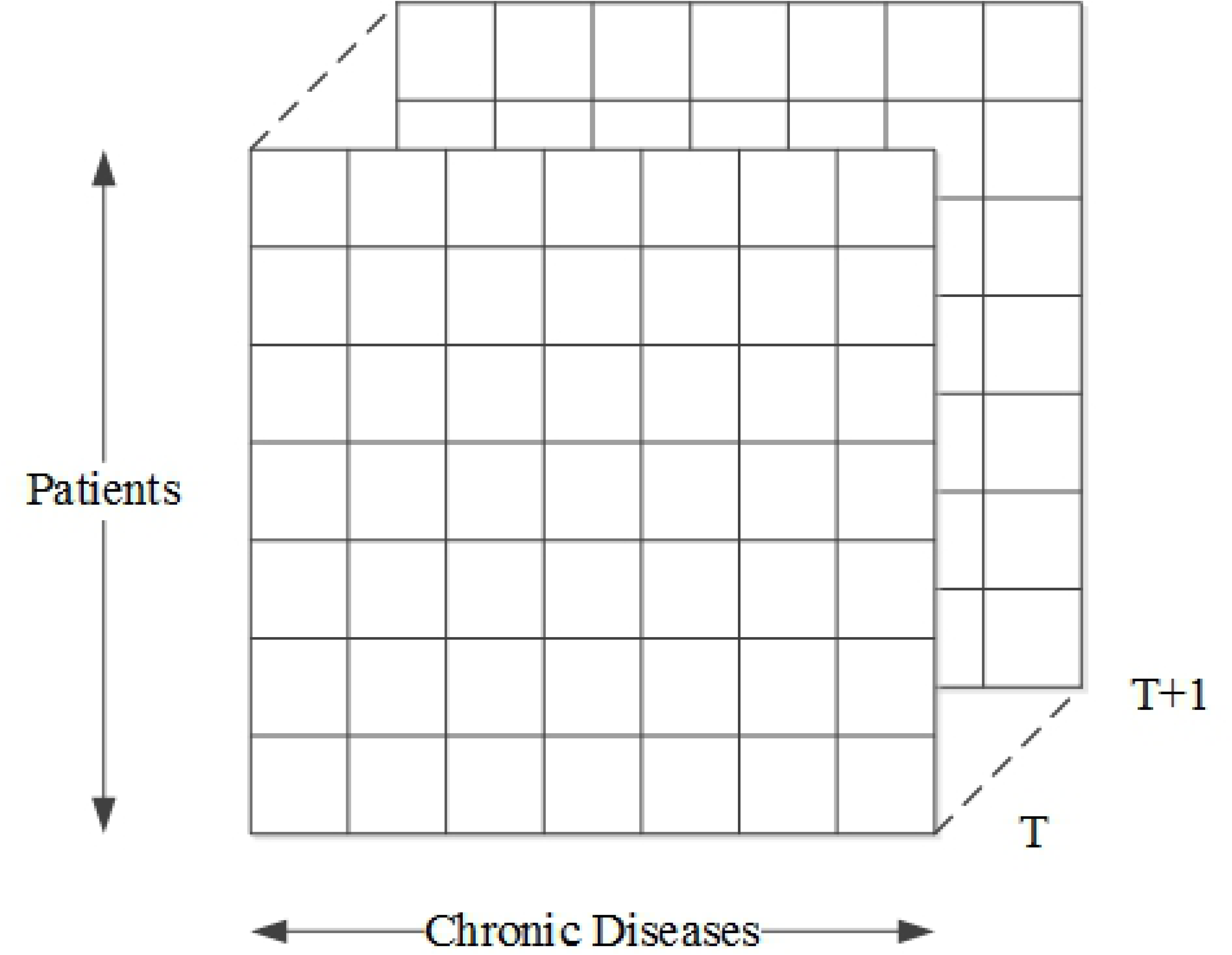
Illustration of tensor representation of the ternary interrelations of patients, diseases and sequence.

We perform NTF on the observed tensor ***χ***. Subsequently, the risk of chronic disease *j* for patient *i* in the future can be represented as the inner product of extracted latent feature vectors in slice *T* + 1 as

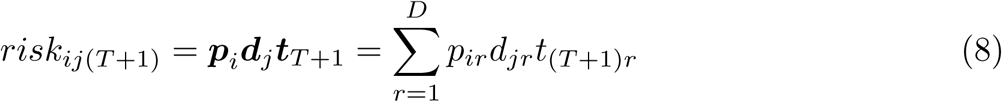

For the illustration of the proposed NTF-based prediction method, hypothetical examples are presented in Fig 4. The size of dots is proportional to value of the corresponding entry. The tensors on the left are observed (thus with discrete count values); the tensors on the right are reconstructed ones by factorizations. The tiny dots on the left represent the non-existence of the corresponding risk factors. After tensor factorizations, the values of entries that were “Nil” in the original tensor are updated in the reconstructed tensor. The updated value of an entry represents the risk of the corresponding disease’s onset for the corresponding patient. For all diseases, the corresponding values in the right tensors can be ranked to determine the risks of different diseases for the individual patients. The rank does not include diseases that already exist in the original tensor (left).

**Fig 4.**
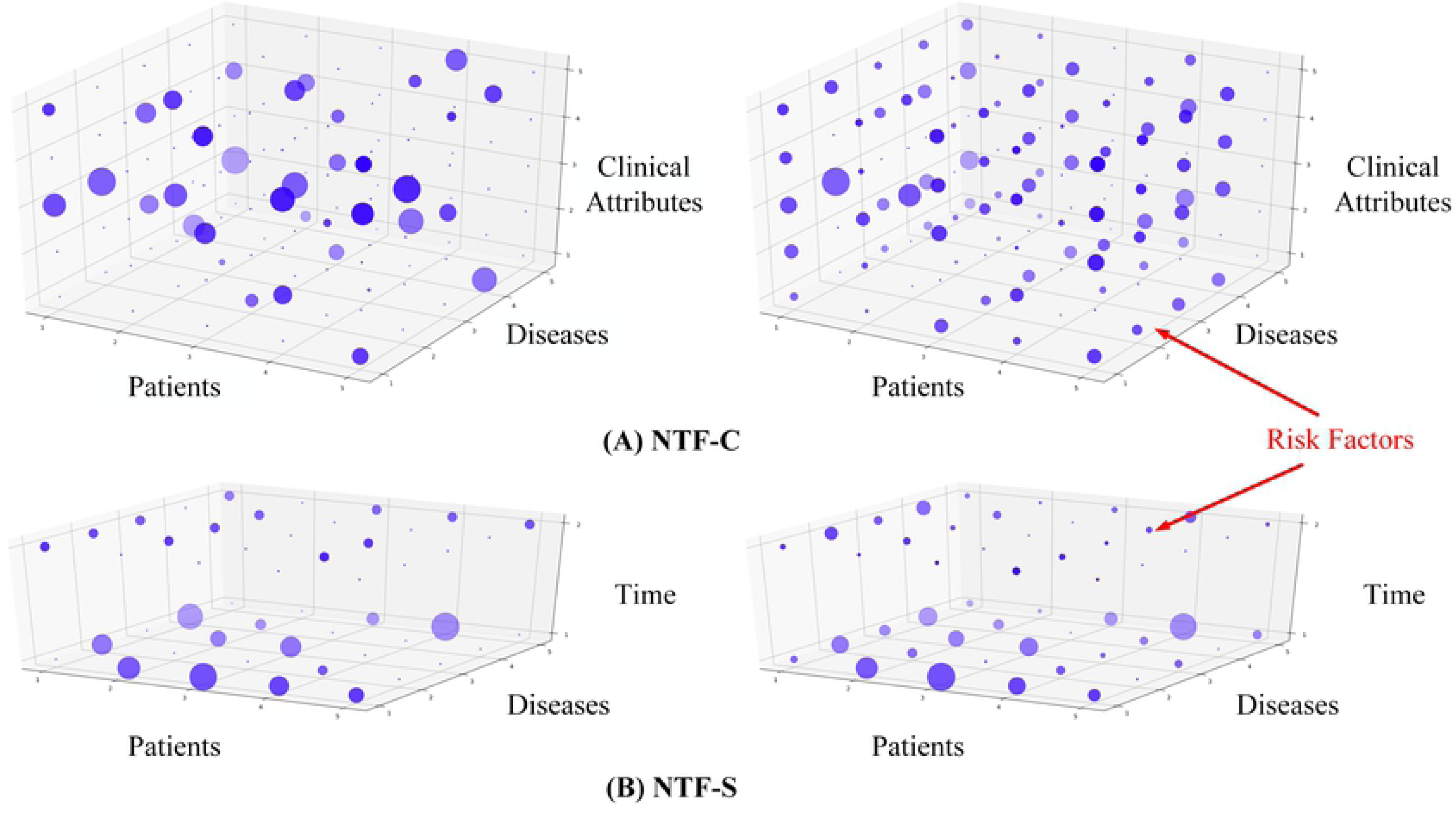
Illustration of the reconstructed tensors (right) of the original tensor (left). (A) The illustration for NTF-C model. (B) The illustration for NTF-S model.

### Individualized Chronic Disease Prediction System

Fig 5 presents the workflow of the proposed approach for chronic disease prediction. First, an appropriate set of data is sampled as the cohort for this study (will be introduced in the next section). Second, the tensor-based on the clinical attributes (for NTF-C) and sequential information (NTF-S) are constructed. Then, we use the sampled EHR data to train the model and evaluate the prediction performance.

**Fig 5.**
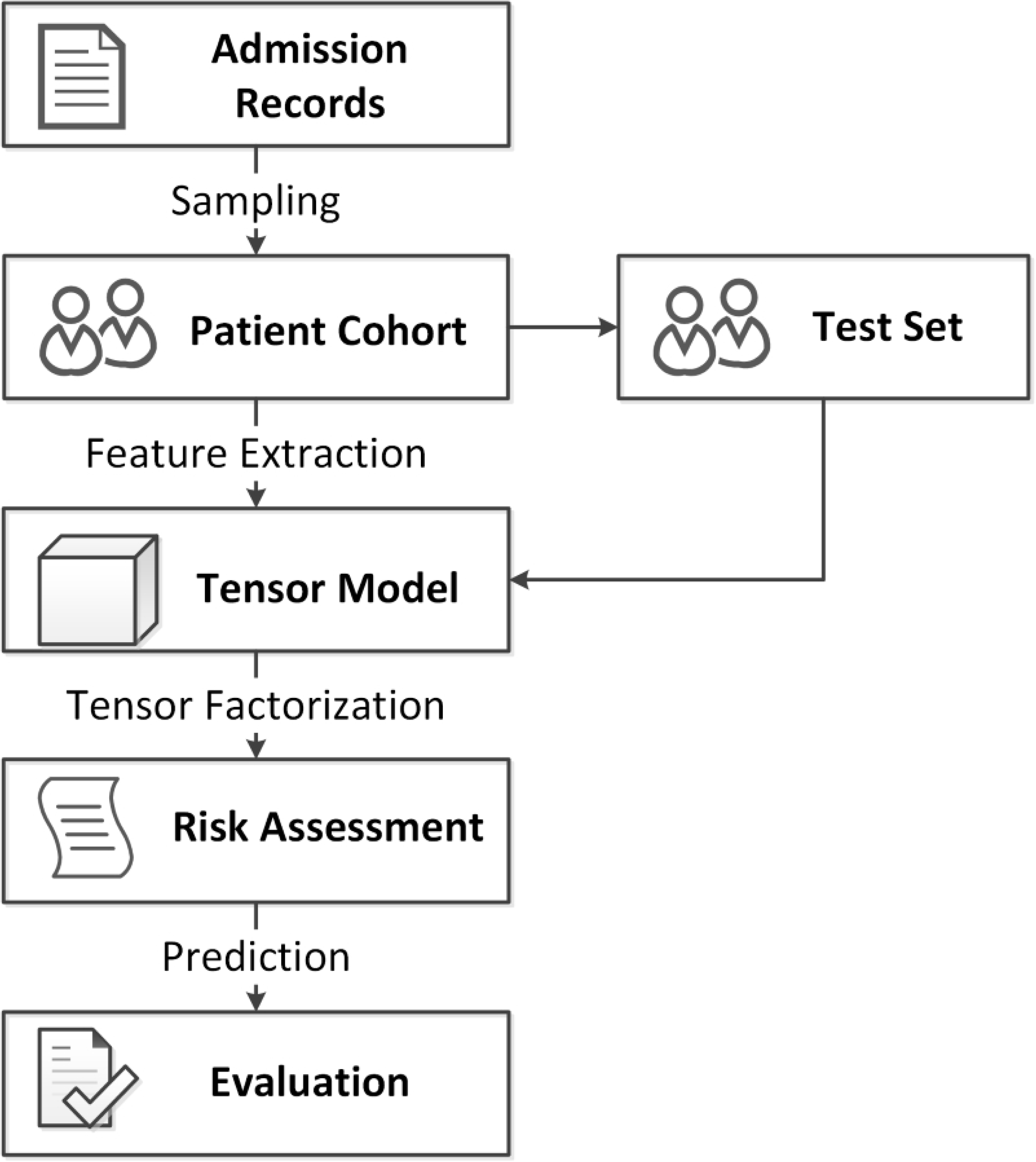
Workflow of the system.

## EXPERIMENTS

To evaluate models for the risk prediction of new chronic diseases, we sampled patients with at least two admission records. The subset of the admission history dataset in 2014 and 2015 with patient assessment information, consisting of 4,168 patients, was used to evaluate the NTF-C method that incorporated the clinical attributes. The subset of admission history dataset (1993-2013), consisting of 5,160 patients (without clinical attributes information), was used to evaluate the NTF-S method that incorporated the sequential patterns.

The 2014-2015 dataset contains clinical attributes including age, blood pressure, pulse, smoking (yes or no), and drinking (yes or no). A total of 690 patients were smokers or ex-smokers, and 381 patients were drinkers or ex-drinkers. Table 2 presents the statistics of other clinical attributes of the patient assessment dataset. To train the model, 2,997 patients with multiple admissions and chronic diseases were set to be the training set, and 1,171 patients who had a new diagnosis in the last admission record were set to be the test set.

**Table 2.** Summary of features from the patient assessments (for 2014-2015 dataset).

|  | mean | std | max | min |
| --- | --- | --- | --- | --- |
| **Number of admissions** | 3.50 | 2.28 | 31 | 2 |
| **Number of diagnoses** | 2.82 | 1.07 | 9 | 2 |
| **Age** | 76.40 | 13.60 | 106 | 19 |
| **BP(systolic)** | 142.19 | 32.30 | 282 | 55 |
| **BP(diastolic)** | 73.05 | 18.66 | 162 | 6 |
| **Pulse** | 81.47 | 22.81 | 196 | 8 |

The 1993-2013 dataset only contains the diagnosis records of patients and could reveal the development of different chronic diseases over time. This dataset was randomly divided into training and test sets for a 5-fold cross-validation.

For the demonstration of the performance of the proposed tensor based methods, 5 benchmark machine learning methods, including logistic regression (LR), multinomial Bayesian classifier (MB), CART decision tree (DT), truncated singular value decomposition (SVD), and the previously introduced NMF, were used to perform the same tasks. In CP factorization, the number of rank-one components, named the rank of tensor, captures the number of potential sub-groups in the tensor. However, determining the value the rank is difficult [5]. Thus, we empirically evaluated different values of the rank and found that the best experiment result could be obtained with rank = 2. The low value of rank is expected due to the sparse nature of the EHR data. The widely used top-k recall was adopted as the evaluation metric. For each patient, the risk score is calculated following Eq (7) or Eq (8) for each potential chronic disease, and then those diseases are sorted in descending order of the score.

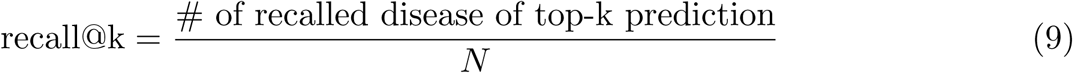

where *N* is the number of future diagnosed diseases in the dataset.

When *k* increases, the *top* − *k* recall will also increase. If *k* is equal to the total number of diseases, then the *top* − *k* recall is 100%, as all possible diseases are covered by the “prediction.” Table 3(a) presents the mean *recall*@*k* using the 2014-2015 dataset.

**Table 3.** Experiment results for the 2014-2015 (patient assessment) and 1993-2013 (admission history) datasets.

| (a) Recall (%) of top-k prediction for the 2014-2015 dataset |  |  |  |  |  |  |  |  |  |  |
| --- | --- | --- | --- | --- | --- | --- | --- | --- | --- | --- |
|  | Top 1 | Top 2 | Top 3 | Top 4 | Top 5 | Top 6 | Top 7 | Top 8 | Top 9 | Top 10 |
| LR | 5.1 | 10.8 | 13.9 | 17.1 | 20.8 | 24.6 | 28.2 | 31.3 | 33.8 | 36.2 |
| MB | 6.6 | 16.7 | 24.3 | 31.9 | 37.8 | 43.3 | 47.8 | 49.9 | 53.0 | 55.6 |
| DT | 7.3 | 16.9 | 23.3 | 30.8 | 40.5 | 47.5 | 51.2 | 54.4 | 57.5 | **60.5** |
| SVD | 11.1 | 17.3 | 25.6 | 36.6 | 43.7 | 48.5 | 52.4 | 55.2 | **58.2** | 59.5 |
| NMF | 11.8 | 19.4 | 27.8 | 38.3 | 45.1 | **49.5** | **53.5** | **57.0** | 58.0 | 59.1 |
| NTF-C | **12.6** | **20.0** | **29.1** | **41.2** | **46.0** | 48.7 | 53.2 | 56.1 | **58.2** | 59.9 |

Table 3(a) also shows the experiment results demonstrating the superior performance of the proposed tensor factorization-based approach.
| (b) Recall (%) of top-k prediction for the 1993-2013 dataset |  |  |  |  |  |  |  |  |  |  |
| --- | --- | --- | --- | --- | --- | --- | --- | --- | --- | --- |
|  | Top 1 | Top 2 | Top 3 | Top 4 | Top 5 | Top 6 | Top 7 | Top 8 | Top 9 | Top 10 |
| LR | 14.4 | 24.1 | 32.5 | 36.7 | 40.5 | 44.8 | 47.5 | 49.5 | 51.1 | 53.5 |
| MB | 14 | 25.9 | 31.3 | 37.3 | 41.5 | 44.2 | 46 | 49.8 | 52.3 | 54.2 |
| DT | 12.7 | 23.9 | 29.1 | 35.1 | 42.5 | 46.7 | 49.1 | 52.4 | 54.1 | 56.5 |
| SVD | 18.3 | 29.8 | 38.2 | 42.4 | 46.9 | 51.5 | 56.6 | 62 | **65.4** | 66.5 |
| NMF | 18.3 | 29.2 | 36.6 | 44.4 | 51.6 | 54.7 | 58 | 61.8 | 62.2 | 64.6 |
| NTF-S | **23.2** | **35.5** | **42.1** | **49.2** | **53.6** | **56.3** | **59.8** | **62.9** | 64.8 | **67.2** |

Table 3(a) also shows the experiment results demonstrating the superior performance of the proposed tensor factorization-based approach. All factorization-based approaches (SVD, NMF and NTF-C) perform better than other benchmark machine learning methods. The tensor factorization-based approach, NTF-C presents the best performance for *top* − 1 to *top* − 5 predictions.

Table 3(b) presents the results for the 1993-2013 dataset (without patient assessment information). The newly proposed NTF-S method consistently outperformed other methods, except for top-9 prediction (only 0.6% lower than SVD). The recall of the top-1 prediction is over 60% higher than the commonly used machine learning methods for risk predictions, such as DT, MB, and LR. The recalls of the NTF-S method for top 5 predictions are higher than 50%. These experiment results demonstrated that modeling the latent high-dimensional associations of ⟨*patient, disease, sequence*⟩ could help us capture useful latent features for prediction.

The performance of NTF-S is better than NTF-C, largely because NTF-S takes advantage of a 23-year patient assessment record dataset that contains rich sequential comorbidity patterns of chronic diseases, such as disease A usually occurred earlier than disease B for one patient. On the other hand, the NTF-C model contains additional clinical attribute information and, thus, performed better than matrix-based models. However, the smaller 2-year patient assessment record dataset limited the performance of the NTF-C model. In our future research, we plan to request a highly comprehensive 23-year EHR dataset (with both sequence and clinical attribute information) to construct an integrated fourth-order ¡patient, disease, clinical attributes, sequence¿ model that takes advantages of both models.

## CONCLUSIONS

To support and inform clinical decision making for healthcare professionals in this aging society, the current study provides an innovative approach to the mapping of an interconnected web of chronic illnesses over the course of aging. With 2 years’ worth of patient assessment records and 23 years’ admission history data from a major hospital in Hong Kong, we demonstrate that tensor factorization is an appropriate approach to capture the latent high-dimensional associations among patients, diseases, clinical attributes, and sequential patterns. The proposed predictive models can predict the onset of new chronic diseases at the individual level with superior accuracy as compared with benchmark machine learning methods.

Benefiting from the capability of tensor to model heterogeneous and multi-aspect data and to extract useful latent features, the proposed tensor factorization-based methods for chronic disease predictive analytics can improve (a) the accuracy of diagnosing underlying and co-occurring chronic conditions and (b) the post-discharge individualized care planning and patient education concerning one’s future risk of chronic and acute conditions given the patients’ chronic conditions at discharge to minimize the likelihood of future hospitalizations. In addition, risk stratification is a proactive strategy to identify at-risk patients with different chronic conditions at discharge into various health management programs, such as telemedicine support with community call centers. In the long run, the implementation of accurate prediction models can prevent readmissions and, thus, reduce the hospital occupancy rate in this aging society.

The study has several limitations. First, the performance of the NTF-C model is dependent heavily on the selection of clinical attributes. However, we are not able to access the typical EHRs data including complete clinical information such as lab test results due to privacy concern without sufficient authorization from the patients. In this research, the six adopted clinical attributes helped in slightly improving the prediction accuracy. These clinical attributes are common risk factors for all chronic diseases and, thus, are not sensitive to specific diseases. How to better select clinical attributes for the model is the focus of our future work. Second, the two proposed third-order tensor-based models incorporate the information of clinical attributes and sequential patterns, respectively. We did not propose a fourth-order tensor to incorporate both information in a single model because of the sparsity problem when we go to the higher order. In our future work, we plan to address this problem by augmenting the observed tensor using the semantic information in knowledge bases. We will also explore other factorization methods, such as coupled tensor factorizations, to solve the sparsity problem. Third, the prediction was made by analyzing the existing EHR data, which could be biased towards the selected cohort, or miss the critical medical information, such as the associations between certain diseases.

## Acknowledgments

This paper is supported by the Theme-Based Research Scheme of the Research Grants Council of Hong Kong under Grant T32-102/14N.

